# Inhibition of PDE5 induces rapid non-genomic increase in hippocampal dendritic spines via specific kinases

**DOI:** 10.1101/2024.12.24.630287

**Authors:** Mika Soma, Mari Ogiue-Ikeda, Shigeo Horie, Minoru Saito, Saria Mabashi, Hirotaka Ishii, Suguru Kawato

## Abstract

Cyclic guanosine monophosphate (cGMP) is a typical neuromodulator that is used in neuronal synapses. Inhibitor of phosphodiesterase 5 (PDE5) could regulates signaling pathways by elevating cGMP levels.

In the current study, we found that the treatments with tadalafil, a PDE5 inhibitor, for 2 h rapidly and non-genomically increased the total density of dendritic spines, using confocal imaging of Lucifer Yellow-injected hippocampal CA1 pyramidal neurons. Since inhibition of PDE5 elevates cGMP levels at synapses, we analyzed downstream nongenomic signaling. We analyzed the involvement of kinase signaling, because rapid postsynaptic modulation often involves kinase networks as observed from many investigations of synaptic functions.

Upon co-treatments of PKG inhibitor (KT5823) with tadalafil, the spine increase was considerably blocked. In addition, co-treatments of GSK-3 inhibitor (I8) also blocked the spine increase induced by tadalafil. However, inhibitors of other representative synaptic kinases, including LIMK, Erk/MAPK, PKA, PKC and PI3K, did not suppress the tadalafil-induced spine increases.

PDE5 inhibitors have attracted attention on their recovery effects from pathological damages, including memory improvement from cognitive impairment by Stroke, Amyloid beta accumulation and tau phosphorylation. The current finding of tadalafil function about rapid modulation of neural plasticity may add new elemental steps of anti-aging capacity against cognitive decline.

## 1 Introduction

Phosphodiesterase 5 (PDE5) is an enzyme that affects cell signaling. Inhibiting PDE5 can relax smooth muscles and increase blood flow to specific areas of the body, which is why PDE5 inhibitors such as tadalafil and sildenafil are used in daily clinical practice for treatment of benign prostatic hyperplasia (BPH), lower urinary tract symptom (LUTS), and erectile dysfunction (ED). Tadalafil regulates signaling pathways by elevating cyclic guanosine monophosphate (cGMP) levels, which decreases in intracellular Ca levels, leading to relaxation of smooth muscles, resulting in improvement of prostatic hyperplasia (Francis et al 2010).

Functions of PDE5 are also investigated in neurons and the brain. PDE5 inhibitors have emerged as a potential therapeutic strategy for neurodegenerative diseases and memory performance. The expression of PDE5 mRNA is observed in the CA1, CA3 and DG regions of mouse and human hippocampus (García-Barroso et al 2013). Not only sildenafil but also tadalafil is observed to cross the blood–brain barrier (García-Barroso et al 2013). In the brain and the hippocampus, the activation of nitric oxide (NO)/cGMP/PKG pathway has an important functional role, via elevation of cGMP levels induced by inhibition of PDE5 (Ko et al 2009) (Zhang et al 2006).

For example, tadalafil elevated cGMP levels and improved short-term memory by suppressing ischemia-induced apoptosis of hippocampal neuronal cells in rodents (Ko et al 2009). Tadalafil improved neurological functional recovery in a rat model of embolic stroke (Zhang et al 2006).

In addition, other PDE5 inhibitors, Zaprinast and Vardenafil, also improved the early memory performance of rodents (Prickaerts et al 2004) (Bollen et al 2014).

PDE5 inhibitors also have attracted much attention on their anti-aging function. For example, tadalafil and sildenafil are useful for suppression of Amyloid β (Aβ) accumulation and tau phosphorylation in Alzheimer’s model animals, in addition to clinical therapy of urological disease. Interestingly, sildenafil is indicated as a best drug for treatments of mild cognitive impairment (MCI) from in silico network analysis (Fang et al 2021).

Both tadalafil and sildenafil have been reported to improve cognitive function in Alzheimer’s disease (AD) mouse models (García-Barroso et al 2013, Puzzo et al 2009). PDE5 is upregulated in AD brains with a decrease in cGMP in the cerebrospinal fluid (CSF) in AD patients (Cuadrado-Tejedor et al 2017), drawing attention to the possible therapeutic potential of PDE5 inhibitors in AD (García-Barroso et al 2013, Puzzo et al 2009). It has been proposed that the activation of the cGMP-responsive element-binding protein (CREB) pathway, which is crucial for memory formation, may contribute to the amelioration of AD symptoms observed with PDE5 inhibitors in animal model (García-Osta et al 2012, Puzzo et al 2005).

Chronic treatments with sildenafil reverses the memory deficits in two different transgenic mouse models of AD (Tg 2576 and APP/PS1), suggesting that PDE5 may be a target for the treatment of these neurodegenerative diseases (Cuadrado-Tejedor et al 2011, Puzzo et al 2009).

There are more than several reports on the chronic effects of PDE5 inhibitors on the brain, but much less investigations are reported on the acute effects.

In wild-type litter mice, brief exposure to oligomeric Aβ_1-42_ (20 min exposure) induced impairment in LTP at CA1 region of the hippocampal slices (Puzzo et al 2005). Interestingly, cGMP analogs (5-10 min perfusion) counteracted the Aβ-induced impairment in LTP and induced CREB phosphorylation (Puzzo et al 2005), suggesting that upregulation of the cGMP pathway may be protective in AD. On the other hand, in APP/PS1 AD model mice, brief perfusion (for 10 min) of sildenafil to hippocampal slices reversed the impairment of LTP in 3 month-old APP/PS1 mice (Puzzo et al 2009). Interestingly such rapid effects in LTP by sildenafil was not observed in wild-type litter mice.

The aim of the current work is to reveal an accurate signaling pathway by analyzing dendritic spine imaging. We found how tadalafil rapidly modulates synaptic plasticity via cGMP/ protein kinase G (PKG)/glycogen synthase kinase 3β (GSK-3β)/ cofilin pathway in dendritic spines of the rat hippocampus. This could be one of the initial step in recovery of structural impairment of synapses in neuronal diseases, including MCI. Interestingly sildenafil is indicated as a best drug for treatments of MCI using in silico network analysis (Fang et al 2021). Therefore, the new finding of tadalafil-induced rapid modulation of neural plasticity may add another impact on anti-aging capacity against early stage of MCI.

## 2 Materials and Methods

### 2.1 Animals

Young adult male Wistar rats (12 week-old, 320-360 g) were purchased from Tokyo Experimental Animals Supply (Japan). All animals were maintained under a 12 h light/12 dark cycle and free access to food and water. The experimental procedure of this research was approved by the Committee for Animal Research of Teikyo University.

### 2.2 Chemicals

Tadalafil, inhibitor Ⅷ (I8, GSK-3β inhibitor), H89 (PKA inhibitor), chelerythrine (PKC inhibitor), LIMKi (LIMK inhibitor) and LY294000 (PI3K inhibitor) were purchased from Sigma-Aldrich (USA). U0126 was purchased from Fujifilm Wako Pure Chemicals (Japan). KT5823 (PKG inhibitor) was purchased from Cayman Chemical (USA). Lucifer Yellow was obtained from Molecular Probes (USA).

### 2.3 Slice preparation

Adult male rats were deeply anaesthetized by isoflurane and decapitated. Immediately after decapitation, the brain was removed from the skull and placed in ice-cold oxygenated (95% O_2_, 5% CO_2_) artificial cerebrospinal fluid (ACSF) containing (in mM): 124 NaCl, 5 KCl, 1.25 NaH_2_PO_4_, 2 MgSO_4_, 2 CaCl_2_, 22 NaHCO_3_, and 10 D-glucose (all from Wako); pH was set at 7.4. The dorsal hippocampus was then dissected and 400 µm thick transverse slices to the long axis, from the middle third of the dorsal hippocampus, were prepared with a vibratome (Dosaka, Japan). These slices were ‘fresh’ slices without ACSF incubation. Slices were then incubated in oxygenated ACSF for 2 h (slice recovery processes) in order to obtain widely used ‘acute slices’.

### 2.4 Imaging and analysis of dendritic spine density and morphology

#### Drug treatments and current injection of Lucifer Yellow

The ‘acute’ slices (used worldwide) were incubated for 2 h with 10 μM tadalafil together with kinase inhibitors. Slices were then fixed with 4 % paraformaldehyde in PBS at 4 °C overnight. Neurons within slices were visualized by an injection of Lucifer Yellow (Molecular Probes, USA) under Nikon E600FN microscope (Japan) equipped with a C2400-79H infrared camera (Hamamatsu Photonics, Japan) and with a 40x water immersion lens (Nikon, Japan).

Current injection was performed with glass electrode filled with 4 % Lucifer Yellow for 2 min, using Axopatch 200B (Axon Instruments, USA). Approximately five neurons within a depth of 10 - 30 µm from the surface of a slice were injected with Lucifer Yellow (Duan et al 2002).

#### Confocal laser microscopic imaging and analysis

The imaging was performed from sequential z-series scans with super-resolution confocal microscope (Zeiss LSM880; Carl Zeiss, Germany) using Airy Scan Mode, at high zoom (× 3.0) with a 63 × oil immersion lens, NA 1.45. For Lucifer Yellow, the excitation and emission wavelengths were 458 nm and 515 nm, respectively. For analysis of spines, three-dimensional image was reconstructed from approximately 30 sequential z-series sections of every 0.45 µm. The applied zoom factor (× 3.0) yielded 23 pixels per 1 µm. The confocal lateral resolution was approximately 0.14 µm. The z-axis resolution was approximately 0.33 µm. Our resolution limits were regarded to be sufficient to allow the determination of the head diameter of spines in addition to the density of spines. Confocal images were deconvoluted with the measured point spread function using Processing Mode of LSM880.

The density of spines as well as the head diameter were analyzed with Spiso-3D (automated software calculating mathematically 3D-geometrical parameters of spines) developed by Bioinformatics Project of Kawato’s group (Mukai et al 2011) (Ooishi et al 2012). Spine image analysis has already often been applied to elucidate effects of neuro-estrogen and neuro-androgen on the hippocampal synapses (Kato et al 2013) (Ikeda et al 2015) (Soma et al 2018) (Murakami et al 2018) (Kumar et al 2020). Spiso-3D has almost an equivalent capacity with Neurolucida (MicroBrightField, USA), furthermore, Spiso-3D considerably reduces human errors and experimenter labor. The single apical dendrite was analyzed separately. The spine density was calculated from the number of spines along secondary dendrites having a total length of 40-60 µm. These dendrites were present within the stratum radiatum, between 100-200 µm from the soma. Spine shapes were classified into three categories as follows. (1) A small-head spine, whose head diameter is smaller than 0.4 µm. (2) A middle-head spine, which has 0.4-0.5 µm spine head. (3) A large-head spine, whose head diameter is larger than 0.5 µm. These three categories were useful to distinguish different responses upon kinase inhibitor application. Small-, middle-, and large-head spines probably have different number of α-amino-3-hydroxy-5-methyl-4-isoxazolepropionic acid (AMPA) receptors, and therefore these three types of spines might have different efficiency in memory storage. The number of AMPA receptors (including GluR1 subunits) in the spine increases as the size of postsynapse increases, whereas the number of N-methyl-D-aspartate (NMDA) receptors (including NR2B subunits) might be relatively constant (Shinohara et al 2008). Because the majority of spines (> 93∼95 %) had a distinct head, and stubby spines and filopodia did not contribute much to overall changes, we analyzed spines having a distinct head.

### 2.5 Statistical analysis

Drug-treated dendrite images were used for spine analysis, and typical images were shown in Fig. 1A. Each dendrite has approx. 50 μm in length including approx. 50 spines. For statistical analysis, we employed two-way ANOVA, followed by Tukey-Kramer multiple comparison’s test. For each drug application analysis, we used approximately 80 dendrites with 4000-5500 spines obtained from 3 rats, 10 slices, 50 neurons. For control dendrite analysis without drug application, we used 50 dendrites with approx. 2500 spines from 3 rats, 10 slices, 50 neurons.

**Figure 1.**
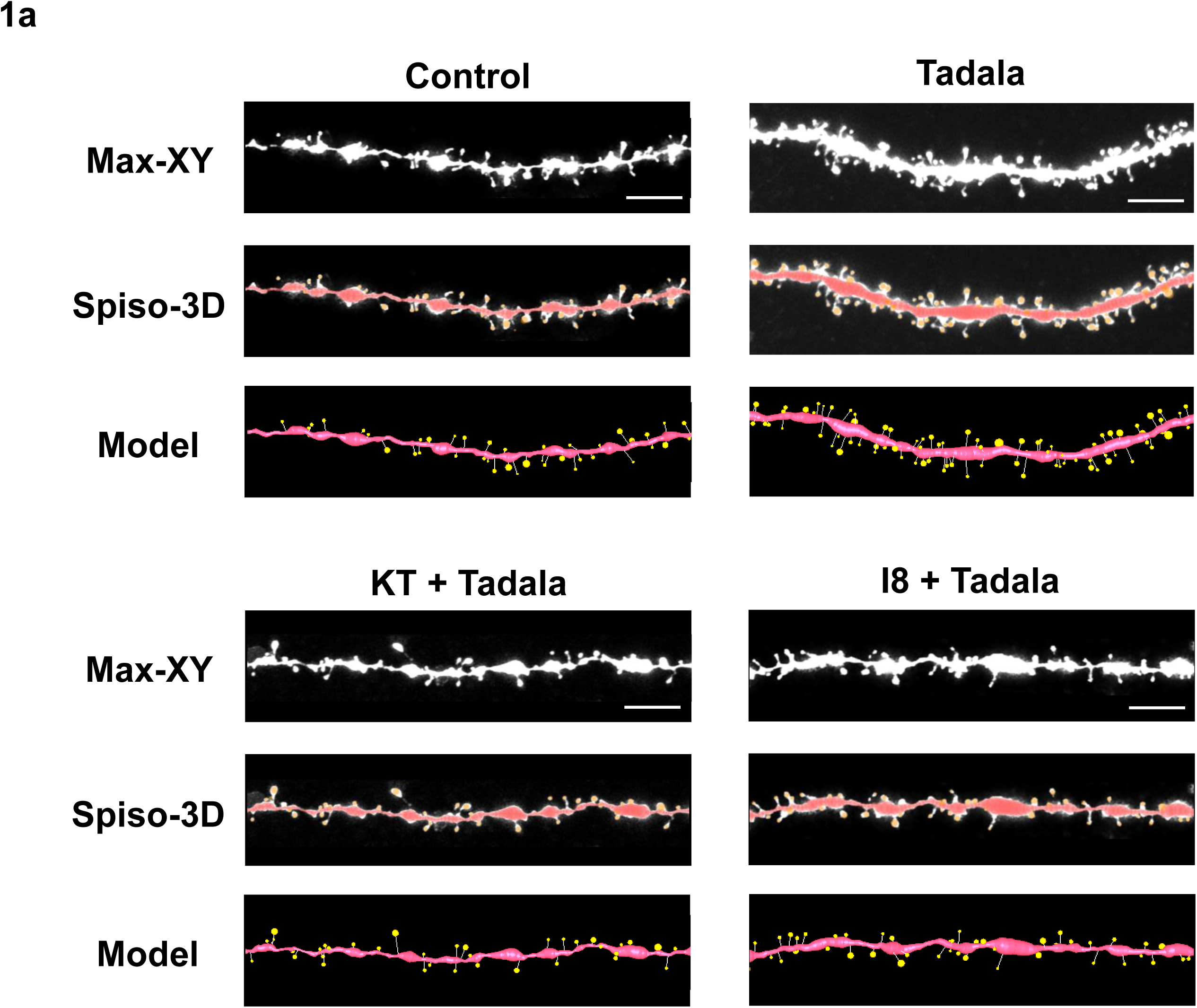
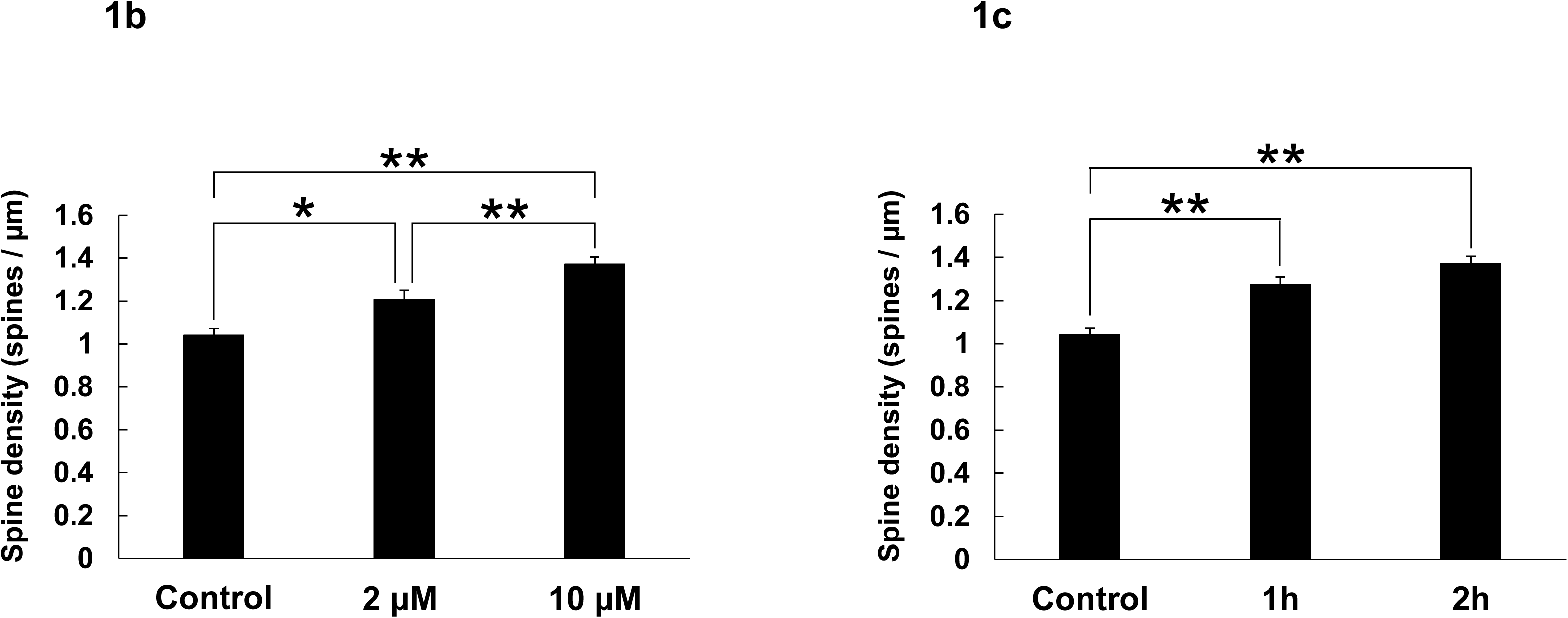
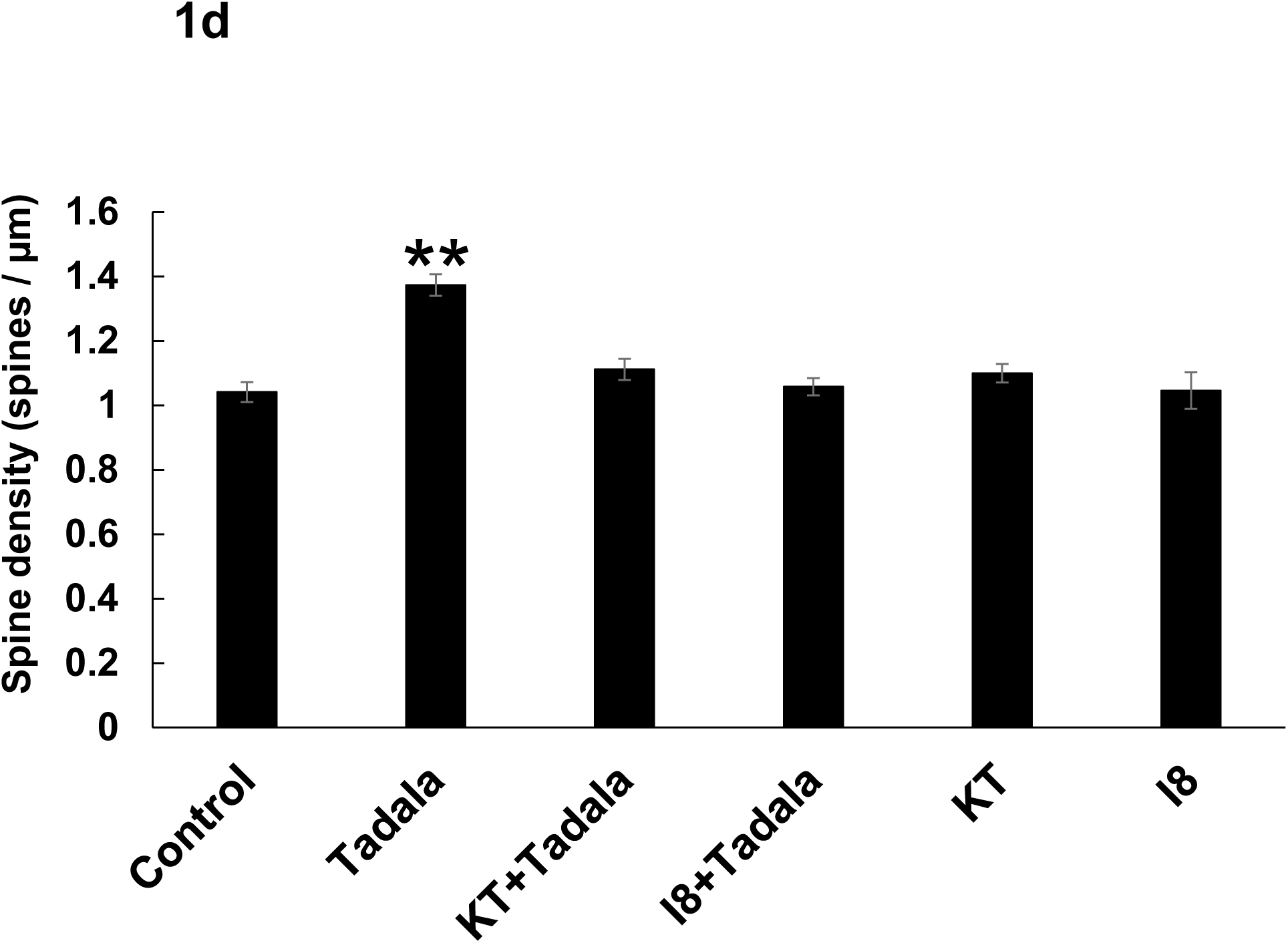
Image analysis demonstrating the changes of spines upon treatments with tadalafil, blockers of PKG and blockers of GSK-3β in adult hippocampal CA1 neurons. **(a)** Spines were analyzed along the secondary dendrites of pyramidal neurons in the stratum radiatum of CA1 neurons. Dendrite without drug-treatment (Control), dendrite after treatments with 10 μM tadalafil for 2 h (Tadala), dendrite after treatments with 10 μM tadalafil plus 2 µM KT5823 (PKG blocker) for 2 h (KT+ Tadala), dendrite after treatments with 10 μM tadalafil plus 10 µM I8 (GSK-3β blocker) for 2 h (I8+ Tadala). Upper images (Max XY) show maximal intensity projections onto XY plane from z-series confocal micrographs. Middle images (Spiso-3D) show the image of dendrite and spines analyzed with Spiso-3D software. Traced dendrite is shown with connected series of red balls and spines are indicated with yellow balls. Lower images (Model) show 3 dimensional model illustration of (Spiso-3D) image. Bar, 5 μm. **(b)** Dose dependency of tadalafil treatments on the total spine density in CA1 neurons. A 2 h treatment in ACSF without tadalafil (Control), with 2 μM tadalafil (2 μM), and with 10 μM tadalafil (10 μM). Vertical axis is the average number of spines per 1 μm of dendrite. **(c)** The time dependency of tadalafil effects on the total spine density in CA1 neurons, after 0 h treatment (Control), 1 h treatment (1 h), 2 h treatment (2 h), in ACSF with 10 μM tadalafil. **(d)** Effect of treatments with tadalafil, KT5823 and I8 on the total spine density in CA1 neurons. Vertical axis is the average number of spines per 1 µm of dendrite. A 2 h treatment in ACSF without drugs (Control), with 10 μM tadalafil (Tadala), with 10 μM tadalafil plus 2 µM KT5823, (KT+ Tadala), with 10 μM tadalafil plus 10 µM I8 (I8+ Tadala), with 2 µM KT5823 only (KT), and with 10 µM I8 only (I8).

## 3 Results

We investigated the rapid Tadalafil (PDE5 inhibitor) effects on the modulation of spinogenesis. Dendritic spine imaging was performed for Lucifer Yellow-injected glutamatergic neurons in acute hippocampal slices of male rats. We analyzed secondary branches of the apical dendrites located 100-200 µm distant from the pyramidal cell body around the middle of the stratum radiatum of CA1 region.

### Analysis of the total spine density as well as spine head diameter

Not only the total spine density but also the morphological changes in spine head diameter induced after 2 h treatments of drugs were analyzed. Since observing the total spine density cannot describe well the complicated different kinase effects, the changes in spine head diameter distribution were also analyzed. Because the majority of spines (> 93∼95 %) had distinct heads and necks, and stubby spines and filopodia did not contribute much to overall changes (less than 5∼7%), we analyzed spines, having distinct heads. We classified these spines with clear heads into three categories based on their head diameter, e.g. 0.2-0.4 µm as small-head spines, 0.4-0.5 µm as middle-head spines, and larger than 0.5 µm as large-head spines (Fig.2).

**Figure 2.**
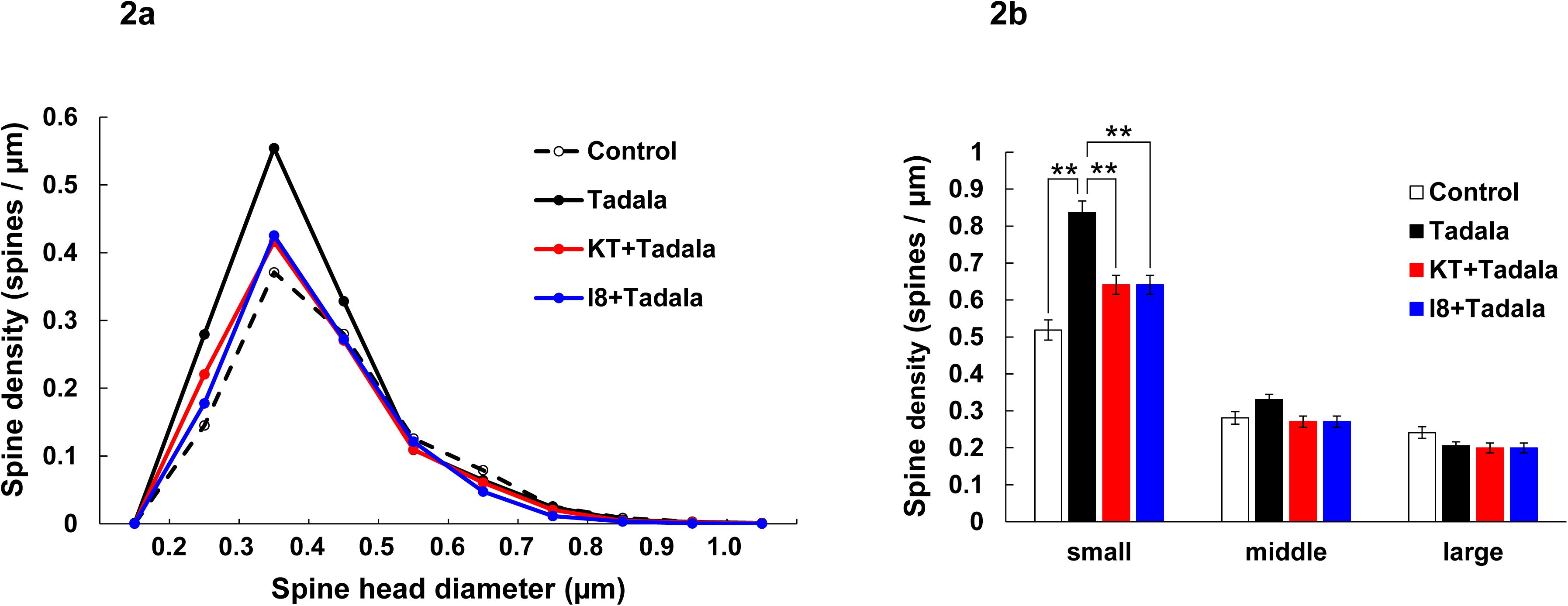
Morphological analysis for effects of tadalafil on spine head diameters in adult hippocampal CA1 neurons. **(a)** Histogram of spine head diameters after a 2 h treatment in ACSF without drugs (Control, black dashed line), with tadalafil (black line), with tadalafil plus KT5823 (red line) and with tadalafil plus I8 (blue line). Spines were classified into three categories depending on their head diameter, e.g. 0.2-0.4 µm as small-head spines, 0.4-0.5 µm as middle-head spines, and larger than 0.5 µm as large-head spines. Vertical axis is the number of spines per 1 µm of dendrite. **(b)** Density of three subtypes of spines. Abbreviations are the same as in (Fig.1D). From left to right, small-head spines (small), middle-head spines (middle), and large-head spines (large) type. ACSF without drugs (Control, open column), with tadalafil (black column), with tadalafil plus KT5823 (red column) and with tadalafil plus I8 (blue column). Vertical axis is the number of spines per 1 µm of dendrite. Results are represented as mean ± SEM. Statistical significance yielded *P < 0.05, **P < 0.01. For 10 μM tadalafil treatments, we investigated, 90 dendrites with approx. 5900 spines from 3 rats, 16 slices and 70 neurons. For control, we used 50 dendrites with approx. 2500 spines from 3 rats, 11 slices and 30 neurons.

Statistical analyses based on classification of the spines into three categories were performed. In control slices (without drug application), the spine density was 0.52 spines/µm for small-head spines, 0.28 spines/µm for middle-head spines, and 0.24 spines/µm for large-head spines (Fig.2). In order to investigate intracellular signaling pathways of kinases involved in the tadalafil-induced spinogenesis, here, we analyzed the contribution of tadalafil and downstream kinases with their selective inhibitors.

### 3.1 Effects of tadalafil and blocking of protein kinase G (PKG) and glycogen synthase kinase 3β (GSK-3β)

#### Dose dependence of tadalafil effects

Dose dependent effects of tadalafil on the total spine density was examined by applying tadalafil at 0 (control), 2 and 10 μM for 2 h (Fig. 1B). Total spine density was significantly increased by applying tadalafil at 2 μM (1.21 spines/µm) and 10 μM (1.37 spines/µm) compared with 0 μM (1.04 spines/µm). [F(2,183)=21.3, p<0.0001, one-way ANOVA, p=0.015 for control vs 2 μM tadalafil, p<0.0001 for control vs 10 μM tadalafil, p=0.005 for 2 μM tadalafil vs 10 μM tadalafil, Tukey-Kramer].

#### Time dependence of tadalafil effects

Time dependent effects of tadalafil on the total spine density was examined by applying tadalafil for 1 h and 2 h at 10 μM (Fig. 1C). Total spine density was significantly higher at 2 h (1.37 spines/µm) than 1 h (1.27 spines/µm). [F(2,200)=21.8, p<0.0001, one-way ANOVA, p<0.0001 for control vs 1 h tadalafil, p<0.0001 for control vs 2 h tadalafil, p=0.0900, for 1 h tadalafil vs 2 h tadalafil, Tukey-Kramer].

From these results, we chose 10 μM tadalafil with 2h treatments, for the following tadalafil-spine experiments.

#### Total spine density analysis

The treatments with tadalafil induced significant changes in the total spine density as analyzed by two-way ANOVA followed by a Tukey-Kramer multiple comparison’s test. A 2h treatment with 10 μM tadalafil, the total spine density was significantly increased to 1.37 spines/µm from the control density of 1.04 spines/µm (Fig. 1D). This tadalafil-induced increase in the total spine density was suppressed by blocking of PKG through co-incubation of 10 μM tadalafil and 2 µM KT5823 (PKG inhibitor). [F(1,285)=22.94, p<0.0001, two-way ANOVA, p<0.0001, for control vs tadalafil, p<0.0001, for tadalafil vs KT5823+ tadalafil, Tukey-Kramer].

The tadalafil-induced increase in the total spine density was also suppressed by blocking of GSK-3β through co-incubation of 10 μM tadalafil and 10 µM I8 (GSK-3β inhibitor) (Fig. 1D). [F(1,221)=16.83, p=0.0001 two-way ANOVA, p < 0.0001, for control vs tadalafil, p<0.0001, for tadalafil vs I8+ tadalafil, Tukey-Kramer].

Note that treatments with only KT5823 and only I8, individually, did not significantly affect the total spine density within experimental error (Fig. 1D). [p=0.22, for control vs KT5823, p=0.93 for control vs I8, Tukey-Kramer]. Therefore, the observed inhibitory effects are not due to simple inhibitors effects.

#### Morphology analysis of spines

From spine head diameter analysis, after 2 h treatments with 10 μM tadalafil, the density of small-head spines significantly/considerably increased [p < 0.0001, t-test] (Figs. 2A, 2B). The density of middle-head spines increased moderately and large-head spines did not increase [p=0.041, for middle-head, p=0.058, for large-head, t-test] (Figs. 2A, 2B).

Blocking PKG by KT5823 and GSK-3β by I8 considerably reduced the tadalafil effects on the dendritic spine densities, by decreasing the density of the small-head spines. [F(1,285)=15.92, p=0.0001, two-way ANOVA, p < 0.0001, for tadalafil vs KT5823+tadalafil, F(1,221)=31.01, p<0.0001 two-way ANOVA, p < 0.0001, for tadalafil vs I8+tadalafil, Tukey-Kramer]. Significant changes in the middle-head and large-head spines did not occur. [for KT5823, F(1,285)=11.03, p=0.001, two-way ANOVA, p=0.034, for tadalafil vs KT5823+tadalafil, for middle-head, F(1,285)=0.347, p=0.556, two-way ANOVA, for large-head] [for I8, F(1,221)=0.361, p=0.549 two-way ANOVA, for middle-head, F(1,221)=6.685, p=0.010 two-way ANOVA, p=0.50, for tadalafil vs I8+tadalafil, Tukey-Kramer, for large-head].

### 3.2 Blocking effects of other essential kinases on tadalafil-induced spine increase

Possible involvement of other kinases, including Erk/MAPK, PKA, PKC, LIMK and PI3K in tadalafil-signaling was also investigated (Fig. 3). The tadalafil-induces increase in the total spine density was not suppressed by blocking of MAPK, PKA, PKC, LIMK and PI3K through co-incubation of 10 μM tadalafil with 10 µM U0126 (MAPK inhibitor), 10 µM H89 (PKA inhibitor), 5 µM Chelerythrine (PKC inhibitor), 10 µM LIMKi (LIMK inhibitor) and 10 µM LY294000 (PI3K inhibitor). [p=0.89, for tadalafil vs U0126+tadalafil, p=0.19, for tadalafil vs H89+tadalafil, p=0.09, for tadalafil vs chelerythrine+tadalafil, p=0.10, for tadalafil vs LIMKi+tadalafil, p=0.36, for tadalafil vs LY294000+tadalafil, Tukey-Kramer] [F(1,225)=0.0001, p=0.99, for U0126, F(1,216)=0.2435, p=0.62, for H89, F(1, 206)=0.0004, p=0.98, for chelerythrine, F(1,216)=0.2728, p=0.60, for LIMKi, F(1,234)=0.2123, p=0.6454, forLY294000, two-way ANOVA].

**Figure 3.**
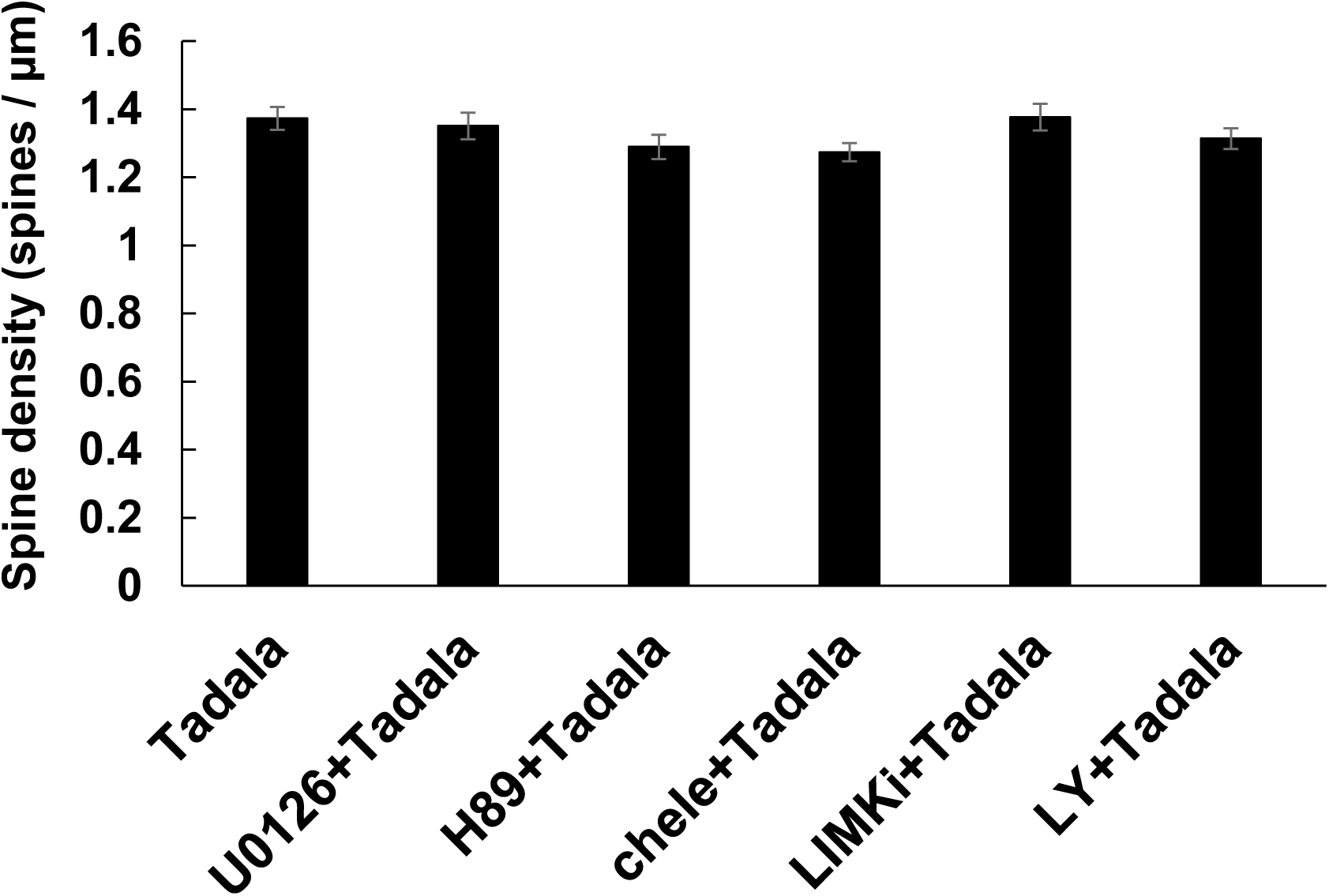
Effect of treatments with tadalafil and several essential kinase blockers on tadalafil-induced spine increase in hippocampal CA1 neurons. Spines were analyzed along the secondary dendrites of pyramidal neurons in the stratum radiatum of CA1 neurons as in Fig.1. Vertical axis is the average numbers of spines per 1μm of dendrite. A 2h treatment in ACSF with 10 μM tadalafil (Tadalafil), with 10 μM tadalafil plus 10 μM U0126 (Erk MAPK inhibitor) (U0126+Tadala), with 10 μM tadalafil plus 10 μM H89 (PKA inhibitor) (H89 +Tadala), with 10 μM tadalafil plus 5 μM chelerythrine (PKC inhibitor) (Chele+Tadala), with 10uM tadalafil plus 10 μM LIMKi (LIMK inhibitor) (LIMKi+Tadala) and with 10 μM tadalafil plus 10 μM LY294002 (PI3K inhibitor) (LY+Tadala). Results are represented as mean ± SEM. Statistical significance yielded *P < 0.05, **P < 0.01. For each condition, we investigated 3 rats, 9 – 12 slices, 47 – 56 neurons, 57 – 75 dendrites and 3200 – 4800 spines.

Note that these kinase inhibitors alone did not significantly affect the total spine density within experimental error, indicating that the observed inhibitory effects are not due to simple inhibitors effects (Hasegawa et al 2015).

## 4 Discussion

PDE5 inhibitors have attracted much attention on their anti-aging function. For example, tadalafil and sildenafil are useful for suppression of Amyloid*β*accumulation and tau phosphorylation in Alzheimer’s model animals, in addition to clinical therapy of urological disease. Interestingly, sildenafil is indicated as a best drug for treatments of mild cognitive impairment (MCI) from in silico network analysis (Fang et al 2021). Therefore, the new finding of tadalafil’s function about rapid modulation of neural plasticity may add another impact on anti-aging capacity against cognitive decline.

In the current study, we found a new tadalafil-dependent rapid spine signaling including PKG with cofilin pathway, as well as GSK-3β (glycogen synthase kinase–3β) with cofilin pathway. Both PKG and GSK-3β signal pathway may induce actin polymerization and spine increase as illustrated in Figure 4 model.

**Figure 4.**
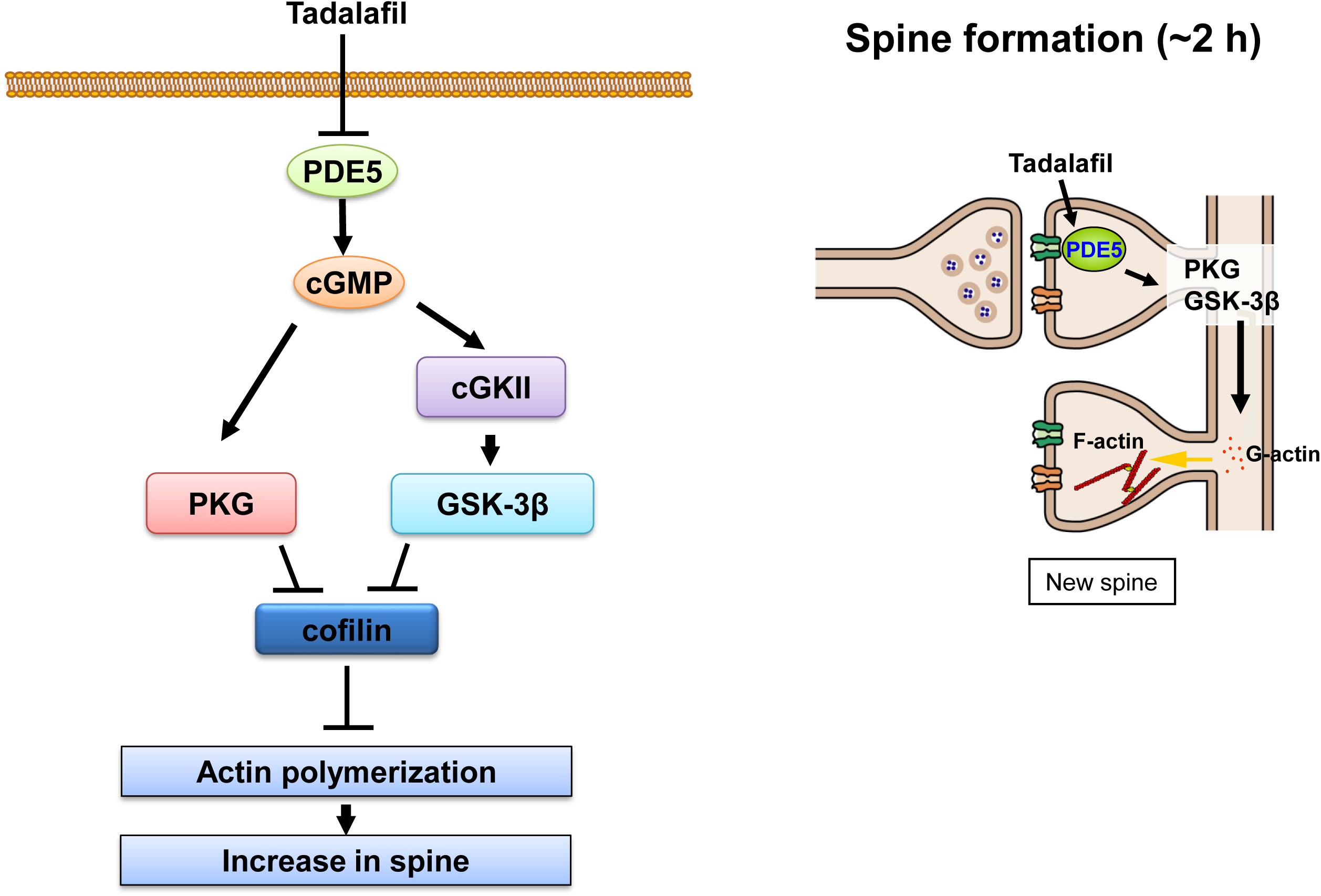
Model illustration of tadalafil-mediated signaling in non-genomic effects on spinogenesis. Tadalafil inhibits PDE5 at synapses. cGMP level is then upregulated at spines, leading to PKG activation, inhibition of cofilin, resulting in spine increase. In parallel, after cGMP level is upregulated, CGKII is activated and then GSK-3β is activated, leading to inhibition of cofilin, resulting in spine increase.

Interestingly, other essential kinases, including MAPK, PKA, PKC, PI3K and LIMK, were not involved at all in tadalafil-induced spinogenesis, although these kinases are often involved in non-genomic processes that are modulated by neurotrophic factors such as IGF-1, neuroestrogen and neuroandrogen (Hakuno & Takahashi 2018) (Hasegawa et al 2015) (Hatanaka et al 2015).

### PKG pathway of PDE5 signal

Current investigations of spine modulation revealed that tadalafil could induce rapid increase of spines by actin polymerization via cGMP/PKG/cofilin pathway. This pathway is deduced from cell biological investigations as follows: PKG-induced cofilin phosphorylation was indicated from the observation that the increase in PKG expression and PKG activity decreased cofilin modulation, which changed actin dynamics in cell adhesion of pulmonary vascular smooth muscle cells. (Negash et al 2009).

### GSK-3β pathway of PDE5 signal

It is also revealed that tadalafil could induce rapid increase of spines by actin polymerization via cGMP/GSK-3β/cofilin pathway. This pathway is deduced from cell biological investigations as follows: cGMP phosphorylates cGMP-dependent GSK-3β (cGKII), leading to phosphorylation/activation of GSK-3β, resulting in cofilin modulation (Tang et al 2011). This was indicated from the observation with Western blotting analysis of p-GSK-3β and p-cofilin in mouse neutrophils (Tang et al 2011) and in ATDC5 cells (Kawasaki et al 2008). GSK-3β signal pathway might also induce microtubule stabilization via tau phosphorylation.

### Little contribution of PDE11 inhibition by tadalafil in dorsal hippocampus

Tadalafil can inhibit not only PDE5 but also PDE11A. However, PDE11 is expressed primarily in the ventral hippocampus and the PDE11 expression level in the dorsal hippocampus is probably less than 5% of that in the ventral hippocampus as judged by immunostaining (Hegde et al 2016) (Kelly et al 2014). In addition, tadalafil inhibits PDE5 much stronger than PDE11 as judged from the specificity index EC50 (EC50= 9.4 (PDE5) and 67 nmol/L (PDE11) (Weed et al 2015). Therefore, the tadalafil-induced spine increase was mainly induced by PDE5 inhibition, because we used the dorsal hippocampus in the present study.

### Chronic effects

PDE5 inhibitors, including tadalafil, are shown to activate CREB pathway, therefore chronic slow effects have been intensively investigated (García-Barroso et al 2013) (Cuadrado-Tejedor et al 2017). For example, PDE5 inhibitors have emerged as a potential therapeutic strategy for neurodegenerative, and memory loss diseases.

Many previous investigations focus on chronic effects of tadalafil and sildenafil (e.g., 4 −10 weeks treatments) on the hippocampal neurons of AD model mice (J20, APP/PS1) (García-Barroso et al 2013) (Cuadrado-Tejedor et al 2017). In chronic regulation, signaling pathway of cGMP/PKG/CREB was indicated. This pathway may regulate slow genomic processes upon binding of p-CREB to CRE binding site on genes, leading to gene transcription and protein synthesis related to neuronal plasticity, resulting in improvement of damaged synaptic plasticity including LTP and spine modulation (García-Barroso et al 2013) (Cuadrado-Tejedor et al 2015) (Cuadrado-Tejedor et al 2017).

Concerning AD problems, since almost all clinical trials of therapies for AD patients have not succeeded until now, with nearly 99% failure (Cummings et al 2014), it has been highlighted the need for approaches to achieve preventive therapy during MCI states, rather than approaches to decrease hyper-phosphorylated tau and amyloid plaques in serious AD states. For this, activation of nitric oxide (NO)/cGMP/PKG pathway has an important functional role which induces elevation of cGMP by inhibiting PDE5 (Zhang et al 2006) (Ko et al 2009).

Note that, expression of PDE5 was also found in human hippocampus (historically being difficult to detect), not only in rodent hippocampus (García-Barroso et al 2013). PDE5 is upregulated in brains of AD patients, with a decrease in cGMP in the cerebrospinal fluid (CSF) (Ugarte et al 2015).

Recently, PDE5 inhibitor sildenafil was indicated as a best drug for treatments of MCI (Fang et al 2021). In silico network medicine analysis of drugs for amyloid and tau pathology by group of F. Cheng described how their network-based analysis of AD-related genes fished through 1,608 approved drugs for possible connections to amyloid and tau pathology (Fang et al 2021). A top hit was sildenafil, a phosphodiesterase 5 inhibitor that crosses the blood-brain barrier (Fang et al 2021). However, note that, tadalafil was unfortunately not included in these in silico network medicine analyses.

### Future perspective

Since we demonstrated that tadalafil rapidly potentiates synaptic plasticity of hippocampal neurons, the rapid recovery of cognitive function could be found by analysis of the clinical records of the hospital. Not only chronic long term treatments but also repeated short term treatments of tadalafil may also be useful for prevention of age-dependent cognitive decline at MCI stage.

## 5 Acknowledgement

Department of Cognitive Neuroscience, Teikyo Univ. is an endowment department, supported with an unrestricted grant from Japan Tobacco Inc.

## 6 Author Contributions

SK conceived and designed the study. MS, MO-I and SM conducted the experiments and analysis of the data. SK wrote the manuscript. SH, MS and HI contributed to discussions. All authors provided feedback on the manuscript.

## 7 Conflict of Interest Statement

The authors declare no conflict of interest.

## Notes

### Competing Interest Statement

The authors have declared no competing interest.

### Summary of Updates

I corrected line 347-345 in Discussion in order to improve the paragraph [Little contribution of PDE11 inhibition by tadalafil in dorsal hippocampus]. The paragraph [Future perspective, line 390-] was also improved. I added the new author Saria Mabashi, because of her contribition to re-examination in the dendritic spine analysis.

## References

Bollen E, Puzzo D, Rutten K, Privitera L, De Vry J, et al. 2014. Improved long-term memory via enhancing cGMP-PKG signaling requires cAMP-PKA signaling. Neuropsychopharmacology 39: 2497–505

Cuadrado-Tejedor M, Garcia-Barroso C, Sánchez-Arias JA, Rabal O, Pérez-González M, et al. 2017. A First-in-Class Small-Molecule that Acts as a Dual Inhibitor of HDAC and PDE5 and that Rescues Hippocampal Synaptic Impairment in Alzheimer’s Disease Mice. Neuropsychopharmacology 42: 524–39

Cuadrado-Tejedor M, Garcia-Barroso C, Sanzhez-Arias J, Mederos S, Rabal O, et al. 2015. Concomitant histone deacetylase and phosphodiesterase 5 inhibition synergistically prevents the disruption in synaptic plasticity and it reverses cognitive impairment in a mouse model of Alzheimer’s disease. Clin Epigenetics 7: 108

Cuadrado-Tejedor M, Hervias I, Ricobaraza A, Puerta E, Pérez-Roldán JM, et al. 2011. Sildenafil restores cognitive function without affecting β-amyloid burden in a mouse model of Alzheimer’s disease. Br J Pharmacol 164: 2029–41

Cummings JL, Morstorf T, Zhong K. 2014. Alzheimer’s disease drug-development pipeline: few candidates, frequent failures. Alzheimers Res Ther 6: 37

Duan H, Wearne SL, Morrison JH, Hof PR. 2002. Quantitative analysis of the dendritic morphology of corticocortical projection neurons in the macaque monkey association cortex. Neuroscience 114: 349–59

Fang J, Zhang P, Zhou Y, Chiang CW, Tan J, et al. 2021. Endophenotype-based in silico network medicine discovery combined with insurance record data mining identifies sildenafil as a candidate drug for Alzheimer’s disease. Nat Aging 1: 1175–88

Francis SH, Busch JL, Corbin JD, Sibley D. 2010. cGMP-dependent protein kinases and cGMP phosphodiesterases in nitric oxide and cGMP action. Pharmacol Rev 62: 525–63

García-Barroso C, Ricobaraza A, Pascual-Lucas M, Unceta N, Rico AJ, et al. 2013. Tadalafil crosses the blood-brain barrier and reverses cognitive dysfunction in a mouse model of AD. Neuropharmacology 64: 114–23

García-Osta A, Cuadrado-Tejedor M, García-Barroso C, Oyarzábal J, Franco R. 2012. Phosphodiesterases as therapeutic targets for Alzheimer’s disease. ACS Chem Neurosci 3: 832–44

Hakuno F, Takahashi SI. 2018. IGF1 receptor signaling pathways. J Mol Endocrinol 61: T69–T86

Hasegawa Y, Hojo Y, Kojima H, Ikeda M, Hotta K, et al. 2015. Estradiol rapidly modulates synaptic plasticity of hippocampal neurons: Involvement of kinase networks. Brain Res

Hatanaka Y, Hojo Y, Mukai H, Murakami G, Komatsuzaki Y, et al. 2015. Rapid increase of spines by dihydrotestosterone and testosterone in hippocampal neurons: Dependence on synaptic androgen receptor and kinase networks. Brain Res 1621: 121–32

Hegde S, Capell WR, Ibrahim BA, Klett J, Patel NS, et al. 2016. Phosphodiesterase 11A (PDE11A), Enriched in Ventral Hippocampus Neurons, is Required for Consolidation of Social but not Nonsocial Memories in Mice. Neuropsychopharmacology 41: 2920–31

Ikeda M, Hojo Y, Komatsuzaki Y, Okamoto M, Kato A, et al. 2015. Hippocampal spine changes across the sleep-wake cycle: corticosterone and kinases. J Endocrinol 226: M13–27

Kato A, Hojo Y, Higo S, Komatsuzaki Y, Murakami G, et al. 2013. Female hippocampal estrogens have a significant correlation with cyclic fluctuation of hippocampal spines. Front Neural Circuits 7: 149

Kawasaki Y, Kugimiya F, Chikuda H, Kamekura S, Ikeda T, et al. 2008. Phosphorylation of GSK-3beta by cGMP-dependent protein kinase II promotes hypertrophic differentiation of murine chondrocytes. The Journal of clinical investigation 118: 2506–15

Kelly MP, Adamowicz W, Bove S, Hartman AJ, Mariga A, et al. 2014. Select 3’,5’-cyclic nucleotide phosphodiesterases exhibit altered expression in the aged rodent brain. Cell Signal 26: 383–97

Ko IG, Shin MS, Kim BK, Kim SE, Sung YH, et al. 2009. Tadalafil improves short-term memory by suppressing ischemia-induced apoptosis of hippocampal neuronal cells in gerbils. Pharmacology, biochemistry, and behavior 91: 629–35

Kumar D, Koyanagi I, Carrier-Ruiz A, Vergara P, Srinivasan S, et al. 2020. Sparse Activity of Hippocampal Adult-Born Neurons during REM Sleep Is Necessary for Memory Consolidation. Neuron 107: 552–65 e10

Mukai H, Hatanaka Y, Mitsuhashi K, Hojo Y, Komatsuzaki Y, et al. 2011. Automated Analysis of Spines from Confocal Laser Microscopy Images: Application to the Discrimination of Androgen and Estrogen Effects on Spinogenesis. Cereb Cortex

Murakami G, Hojo Y, Kato A, Komatsuzaki Y, Horie S, et al. 2018. Rapid nongenomic modulation by neurosteroids of dendritic spines in the hippocampus: Androgen, oestrogen and corticosteroid. Journal of neuroendocrinology 30

Negash S, Narasimhan SR, Zhou W, Liu J, Wei FL, et al. 2009. Role of cGMP-dependent protein kinase in regulation of pulmonary vascular smooth muscle cell adhesion and migration: effect of hypoxia. Am J Physiol Heart Circ Physiol 297: H304–12

Ooishi Y, Kawato S, Hojo Y, Hatanaka Y, Higo S, et al. 2012. Modulation of synaptic plasticity in the hippocampus by hippocampus-derived estrogen and androgen. J Steroid Biochem Mol Biol 131: 37–51

Prickaerts J, Sik A, van Staveren WC, Koopmans G, Steinbusch HW, et al. 2004. Phosphodiesterase type 5 inhibition improves early memory consolidation of object information. Neurochem Int 45: 915–28

Puzzo D, Staniszewski A, Deng SX, Privitera L, Leznik E, et al. 2009. Phosphodiesterase 5 inhibition improves synaptic function, memory, and amyloid-beta load in an Alzheimer’s disease mouse model. J Neurosci 29: 8075–86

Puzzo D, Vitolo O, Trinchese F, Jacob JP, Palmeri A, Arancio O. 2005. Amyloid-beta peptide inhibits activation of the nitric oxide/cGMP/cAMP-responsive element-binding protein pathway during hippocampal synaptic plasticity. J Neurosci 25: 6887–97

Shinohara Y, Hirase H, Watanabe M, Itakura M, Takahashi M, Shigemoto R. 2008. Left-right asymmetry of the hippocampal synapses with differential subunit allocation of glutamate receptors. Proc Natl Acad Sci U S A 105: 19498–503

Soma M, Kim J, Kato A, Kawato S. 2018. Src Kinase Dependent Rapid Non-genomic Modulation of Hippocampal Spinogenesis Induced by Androgen and Estrogen. Front Neurosci 12: 282

Tang W, Zhang Y, Xu W, Harden TK, Sondek J, et al. 2011. A PLCβ/PI3Kγ-GSK3 signaling pathway regulates cofilin phosphatase slingshot2 and neutrophil polarization and chemotaxis. Dev Cell 21: 1038–50

Ugarte A, Gil-Bea F, García-Barroso C, Cedazo-Minguez Á, Ramírez MJ, et al. 2015. Decreased levels of guanosine 3’, 5’-monophosphate (cGMP) in cerebrospinal fluid (CSF) are associated with cognitive decline and amyloid pathology in Alzheimer’s disease. Neuropathol Appl Neurobiol 41: 471–82

Weed DT, Vella JL, Reis IM, De la Fuente AC, Gomez C, et al. 2015. Tadalafil reduces myeloid-derived suppressor cells and regulatory T cells and promotes tumor immunity in patients with head and neck squamous cell carcinoma. Clin Cancer Res 21: 39–48

Zhang L, Zhang Z, Zhang RL, Cui Y, LaPointe MC, et al. 2006. Tadalafil, a long-acting type 5 phosphodiesterase isoenzyme inhibitor, improves neurological functional recovery in a rat model of embolic stroke. Brain Res 1118: 192–8

